# Spatiotemporal dynamics revealed the dark water community of giant virus from a deep freshwater lake

**DOI:** 10.1101/2024.04.21.590434

**Authors:** Liwen Zhang, Lingjie Meng, Yue Fang, Hiroyuki Ogata, Yusuke Okazaki

## Abstract

Giant viruses significantly regulate the ecological dynamics of diverse ecosystems. Although metagenomics has expanded our understanding of their diversity and ecological roles played in marine environments, little is known about giant viruses of freshwater ecosystems. Most previous studies have employed short-read sequencing and therefore resulted in fragmented genomes, hampering accurate assessment of genetic diversity. We sought to bridge this knowledge gap and overcome previous technical limitations. We subjected spatiotemporal (2 depths × 12 months) samples from Lake Biwa to metagenome-assembled genome reconstruction enhanced by long-read metagenomics. This yielded 294 giant virus metagenome-assembled genomes. Of these, 285 included previously unknown species in five orders of nucleocytoviruses and the first representatives of freshwater mirusviruses, which exhibited marked divergence between freshwater- and marine-derived lineages. Notably, 42 (14.3%) genomes were composed of single contigs with completeness values >90%, demonstrating the good performance of our long-read metagenomic assembly. Giant viruses were partitioned across water depths, with most species specific to either the sunlit epilimnion or the dark hypolimnion. Epilimnion-specific members tended to be opportunistic and to exhibit short and intense abundance peaks, in line with the fact that they regulate the surface algal blooms. During the spring bloom, mirusviruses and members of three nucleocytovirus families were among the most abundant giant viruses. In contrast, hypolimnion-specific ones including algaviruses and mirusviruses were typically more persistent in the hypolimnion throughout the water-stratified period, suggesting that they infect hosts specific to the hypolimnion and play previously unexplored ecological roles in dark water-specific microbial ecosystems.

## Introduction

Giant viruses (GVs) are a significant group within the virosphere, exhibiting remarkable diversity, ubiquity, and abundance across various ecosystems such as oceans, freshwater, and soil [1–6]. In marine ecosystems, they are widespread and distributed across the water column. In contrast, the diversity of freshwater GVs has not been well studied despite indications of high diversity, as revealed by the distribution of their major capsid proteins (MCPs) in freshwater environments [4].

Freshwater lakes, known for their complex seasonal and vertical dynamics, have been the subject of extensive studies in terms of the temporal shifts and vertical stratification of plankton and prokaryote communities [7–10]. These studies have revealed the niche preferences of eukaryotic and prokaryotic microbes across seasons and depths, highlighting the existence of a deep-water specific microbiome. A recent study revealed the dominant GVs associated with spring algal blooms in photic zones [11]. However, the ecological dynamics of GVs in freshwater lakes remain poorly understood and no study has previously addressed the existence of GVs specific to the dark and deep layers of a lake.

We comprehensively analyzed the diversity of GVs in a deep freshwater ecosystem via reconstruction of metagenome-assembled genomes (MAGs) for investigating the GV dynamics across seasons and depths. To achieve this, we combined a spatiotemporal sampling strategy with long-read metagenomic sequencing. This enabled us to capture the GV community dynamics within the ecosystem and overcame the problem posed by fragmented assembly of conventional short-read metagenomes [12]. The generation of more continuous contigs via long-read sequencing aids the identification of a full set of marker genes for MAGs, allowing more accurate quality evaluation and taxonomic assignment. Indeed, an increasing number of studies have used long-read sequencing to generate better genomes of isolated GVs [13–15], but no long-read GV MAG has been generated from metagenomic data. Moreover, we developed a pipeline that detected not only GVs of the phylum *Nucleocytoviricota* but also those of *Mirusviricota*, a newly discovered GV phylum [5]. Previous metagenomic studies have often overlooked mirusviruses given their high genomic novelty and chimeric attributes [5]. Indeed, mirusviruses in the freshwater ecosystems remain to be discovered.

We leveraged previously published short- and long-read metagenomic data from Lake Biwa, a deep oligo-mesotrophic freshwater lake of Japan [16]. The data originally targeted the prokaryotic community (size fraction = 0.2–5 µm) and were collected spatiotemporally. GVs have the same size fraction as prokaryotes [17] but were not investigated in the original study. We reanalyzed these data using a custom pipeline that recovers GV genomes from long-read contigs and bins. This led to the reconstruction of circular or nearly complete GV MAGs, including eight novel representatives of freshwater mirusviruses. Moreover, the spatiotemporal data revealed the dynamics of GV communities and their specific occurrences across the depths.

## Materials and Methods

### Data source

We compiled long-read MAGs and contigs derived in a recent study on Lake Biwa, Japan [16]. Dataset samples were collected monthly from May 2018 to April 2019. Throughout the one-year sampling period, thermal stratification occurred from May to December. During each sampling event, water samples from two depths (5 m for the epilimnion and 65 m for the hypolimnion) were collected (24 samples in total). DNA was extracted from the 0.22–5 μm size fraction of each sample and subjected to short-read (MGI DNBSEQ-G400) and long-read metagenomic sequencing (Oxford Nanopore). The contigs used here were those assembled by Flye v2.8 [18] in the previous study. The long-read contigs were polished using both the long and short reads. The detailed workflow and the relevant parameters are described in the original publication [16].

### Reconstruction of long-read GV MAGs

Many contigs were sufficiently long to be considered nearly complete GV genomes (>50 kb) with GV signals (≥1/7 nucleocytovirus marker genes or the mirusvirus HK97 MCP). The seven nucleocytovirus marker genes were those encoding major capsid protein (MCP), DNA polymerase family B (PolB), transcription initiation factor IIB (TFIIB), DNA topoisomerase II (TopoII), packaging ATPase (A32), DEAD/SNF2-like helicase (SFII), and the poxvirus late transcription factor VLTF3 [19]. In addition to these contigs, we retained all 4,648 bins generated in the original study [16], which together allowed exclusion of prokaryotic genomes with CheckM v1.2.2 [20]. MAGs with CheckM completeness scores higher than 15 as bacteria or 20 as archaea were considered to be prokaryotes and therefore excluded (Fig. S1).

Next, we screened for putative GV MAGs using different methods (see Supplementary Methods for the details) to identify nucleocytoviruses and mirusviruses. For nucleocytoviruses, we employed a weighted scoring method based on the presence of 20 nucleocytovirus core genes [21]. The presence or absence of these 20 core genes in each MAG was determined using the function “hmmsearch” of HMMER3 v3.4 with an E-value of 1×10^-3^ [22]. A cut-off of 5.75 was used to exclude non-GV MAGs and to select putative nucleocytovirus MAGs for further examinations [21]. To detect mirusviruses, we screened for the mirusvirus HK97 MCP gene as this is a unique marker of mirusviruses [5]. A MAG was identified as a mirusvirus if the HK97 MCP gene was detected using “hmmsearch” (bitscore >130).

Following the MAG detection, we removed all cellular contamination (Fig. S2) and then excluded chimeric, low-quality, and fragmented GV MAGs prior to downstream analyses (Fig. S3) (see the Supplementary Methods for the details). Finally, redundant GV MAGs generated by the above processes were removed at an average nucleotide identity (ANI) threshold of 95% with dRep v3.2.2 [23]. The resulting nonredundant GV MAGs were species-level representatives that we termed “Lake Biwa giant virus metagenome-assembled genomes” (LBGVMAGs). Each was assigned a unique four-digit serial number as part of the ID, based on the maximum coverage rank across all 24 samples.

### Quality assessment of long-read GV MAGs

We first assessed the diversity-coverage of our MAGs by determining the proportion of uncaptured nucleocytovirus *polB* sequences. The unique nature of this gene, which is single-copy and universal in nucleocytovirus genomes, allowed us to assess how much of the GV diversity in the lake was captured by our MAGs. We performed a blastn search using blast+ v2.15.0 [24] to align all representatives of clustered *polB* sequences (see the Supplementary Methods for the details) against the contigs of our MAGs with an E-value cut-off of 1×10^-5^ and an alignment identity >96% as it reportedly suggests the GV species boundary [25].

To compare the fragmentation levels of our long-read MAGs and those of the short-read MAGs, we compiled nonredundant quality-controlled (high/medium quality) short-read GV MAGs from the Giant Virus Database (GVDB) [19]. Seqkit v2.5.1 [26] was employed to calculate the number of contigs and the N50 value of each MAG. We also determined the POA90 score of each MAG; this metric evaluates unpolished indel errors in long-read assemblies [16]. The quality assessment details are given in the Supplementary Methods.

### Analyses of the phylogenetic diversity and community dynamics of GV MAGs

To evaluate the novelty of our MAGs, we complied a custom database that integrated the GVDB [19] and 697 nucleocytovirus/mirusvirus MAGs recovered from *Tara Oceans* and lodged in the Global Ocean Eukaryotic Viral database (GOEV) [5]. This custom database followed the taxonomic classification of the GVDB. Next, we used fastANI v1.33 with the default parameters to calculate the pair-wise ANIs between this custom database and our MAGs [27]. The alignments were visualized with DiGAlign [28].

For phylogenetic analysis of the nucleocytoviruses, we used the “ncldv_markersearch.py” script with the “-c” parameter to generate a concatenated alignment of seven genes (encoding PolB, SFII, TFIIB, TopoII, A32, VLTF3, and the DNA-directed RNA polymerase alpha subunit [RNAPL]) [3] from our MAGs and reference genomes. Subsequently, we trimmed the alignment at >90% gaps using TrimAl [29], generated a phylogenetic tree with IQ-TREE v2.2.2.6 [30] using Ultrafast Bootstrap [31] (parameters: -wbt -bb 1000 -m LG+I+G4), and visualized the tree using iTOL [32]. The taxonomy of our MAGs was manually determined based on the topology of the tree following the taxonomic classification of the GVDB.

To infer mirusvirus phylogeny, we generated individual phylogenetic trees of HK97 MCP and heliorhodopsin sequences. We searched against an HMM model built from marine mirusviruses [5] to screen for HK97 MCPs using the hmmsearch (bitscore >130), and another HMM model generated from the custom database described above to screen for heliorhodopsins with an E-value of 1×10^-3^. In addition to sequences detected in our MAGs, sequences from marine mirusviruses were included in the tree as references. For the tree of heliorhosopsins, we additionally included heliorhodopsins from the RefSeq database [33]. Note that we excluded sequences of the marine mirusvirus family M7 from both trees due to the phylogenetic instability introduced by them. Alignments of the HK97 MCPs and heliorhodopsins were generated using MAFFT v7.520 [34] with the “L-INS-i” algorithm and trimmed at gaps >90% with trimAl v1.4. Trees were built using IQ-TREE v2.2.2.6 [30, 31, 35] with the Ultrafast Bootstrap parameters “-B 1000 -alrt 1000”. Model “LG+R8” and “VT+F+R8” were used to generate the trees of HK97 MCPs and heliorhodopsins, respectively [5]. Phylogenetic inferences aside, we further compared the genome contents of mirusviruses from marine and freshwater clades as described in the Supplementary Methods.

To determine the relative abundances and spatiotemporal distributions of LBGVMAGs, the coverage and read per kilobase per million reads (RPKM) of each MAG were calculated based on the mapping of short reads from all samples to LBGVMAGs using CoverM v0.6.1 [36] with parameters “--min-read-percent-identity 0.92 -p bwa-mem2”. The taxonomic composition was based on the RPKMs of each order per sample and visualized using the ggplot2 package [37] in R Studio [38, 39]. The beta diversity between different communities was calculated using the vegdist function (method = “bray”) of the vegan package [40] and used for nonmetric multidimensional scaling (NMDS) analysis. The quantitative analysis that compared the compositional variances between the epilimnion and hypolimnion used the same distance matrix, but we limited our analysis to samples collected during the period of water stratification.

The habitat preference of LBGVMAGs was assessed by an indicator termed “P_epi_”, which is the cumulative RPKM in the epilimnion divided by the cumulative RPKM in both epilimnion and hypolimnion during the water stratification period [16]. When P_epi_ was >0.95 or <0.05, the LBGVMAG was defined as epilimnion- or hypolimnion-specific, respectively. The persistence of each LBGVMAG was defined as the longest consecutive months for which the covered fraction of the LBGVMAG was >50% from short-read mapping [16]. Persistence of epilimnion-/hypolimnion-specific LBGVMAGs was calculated for only epilimnion/hypolimnion samples. Statistical evaluation of differences in the mean persistence between the two groups employed the Welch t-test.

## Results

### High quality long-read assembled GV MAGs

Within the 24 samples, 0.5% to 4.2% (mean = 2.1%) of the short reads were mapped onto the LBGVMAGs (hereafter, GV MAGs or MAGs when there is no ambiguity). The percentage of mapped reads was typically larger for the epilimnion than the hypolimnion (Fig. S4). Using these samples, we successfully reconstructed 293 nonredundant species-level GV MAGs (Table S1). Moreover, the high quality of our GV MAGs was verified by comparison with previously reported short-read MAGs, and via completeness assessment. Our long-read-assembled MAGs yielded a significantly longer (*p*-value = 8.376×10^-16^) and lower number of contigs (*p*-value = 6.818×10^-6^) compared to the short-read MAGs (Fig. S5). Notably, we obtained 42 (14.3%) single-contig MAGs with completeness scores >90%, including two MAGs (0074 and 0028) previously identified as circular [16]. CheckV classified 118 (40%) GV MAGs with completeness values >90% as “high-quality” [41], of which 74 (62.7%) MAGs contained all seven marker genes (Fig. 1A).

**Figure 1.**
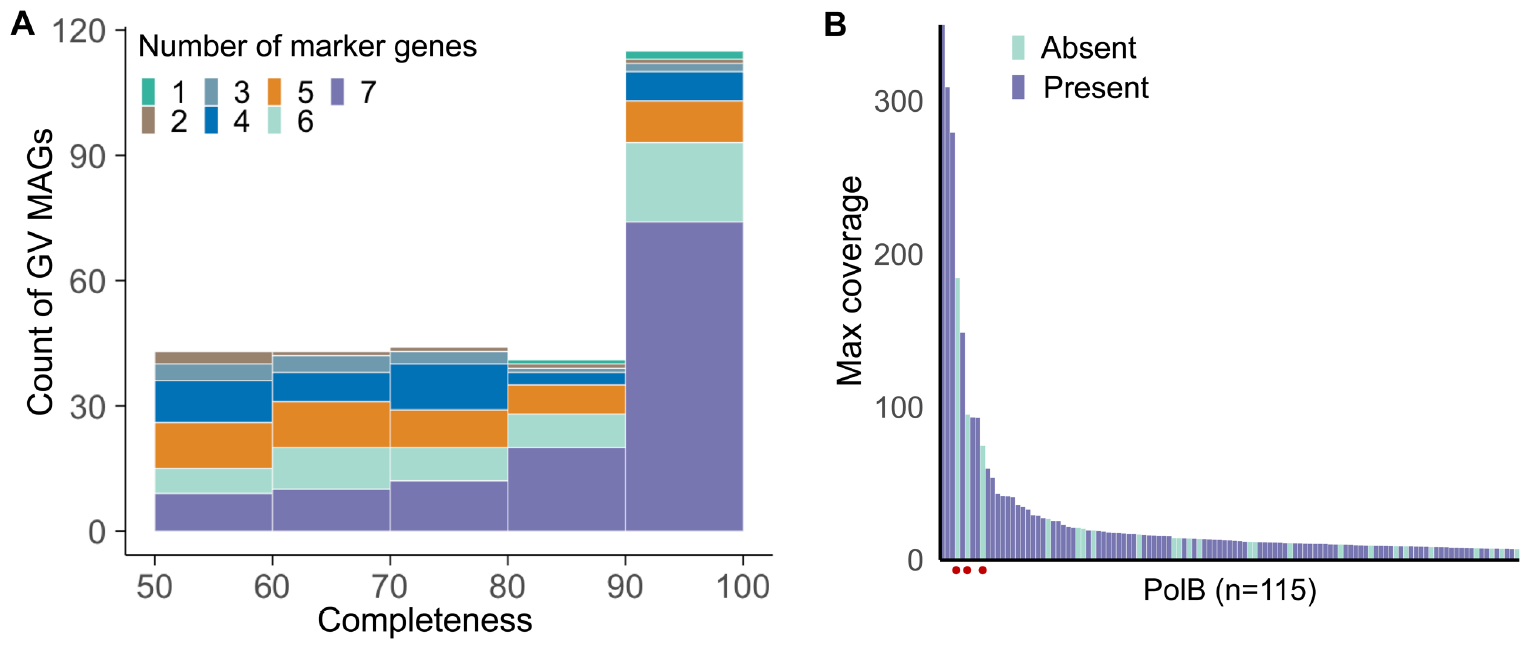
GV MAG quality and diversity coverage. (A) Distribution of the GV MAG completeness scores. Colors indicate the number (maximum: 7) of nucleocytovirus marker genes in each MAG. (B) Presence or absence of abundant *polB* genes in the GV MAGs. “Max coverage” refers to the highest read coverage of the contig from which the *polB* was called among the 24 samples. Red dots indicate the three abundant *polB* genes absent from our MAGs.

A large proportion of the *polB* genes (80%) in the assembled contigs were present in our GV MAGs, indicating that our MAGs represented most of the GV diversity present in the lake (Fig. 1B). Upon closer examination, the three most abundant nucleocytovirus *polB* genes absent from the MAGs were encoded in cellular contigs, suggesting that our pipeline effectively eliminated contamination of cellular sequences. The POA90 score that assessed the performance of contig polishing (see the Supplementary Methods for details) decreased when the short-read coverage was below 7× (Fig. S6), suggesting that a short-read coverage >7× was required for effective nucleotide error correction.

### Expanded diversity of giant viruses

The GV MAGs included mirusviruses (Fig. 2A) and five orders of nucleocytoviruses (Fig. 2B). We identified 285 new species, sharing <95% of their ANIs with known GV genomes (Fig. 2C). Notably, 177 (60.4%) had ANI values <80% (Fig. 2C) or ANI values that could not be calculated because of large divergences from the reference genomes. Of interest, 85.9% of GV MAGs with assigned ANIs were closely related to genomes from freshwater environments (Table S2). We also discovered three GV MAGs of the IM_01 family that were nearly identical (Fig. S7) to the MAGs recovered from Lake Lanier in North America [3, 4]. Specifically, MAGs 0129, 0046, and 0059 exhibited ANIs of 99.2%, 98.2%, and 98.2% to the MAGs from Lake Lanier, respectively. The order *Imitervirales* exhibited pronounced diversity, accounting for 237 (80.9%) MAGs in this study. Also, we recovered 22 *Algavirales*, 16 *Pimascovirales*, eight *Asfuvirales*, and two *Pandoravirales*.

**Figure 2.**
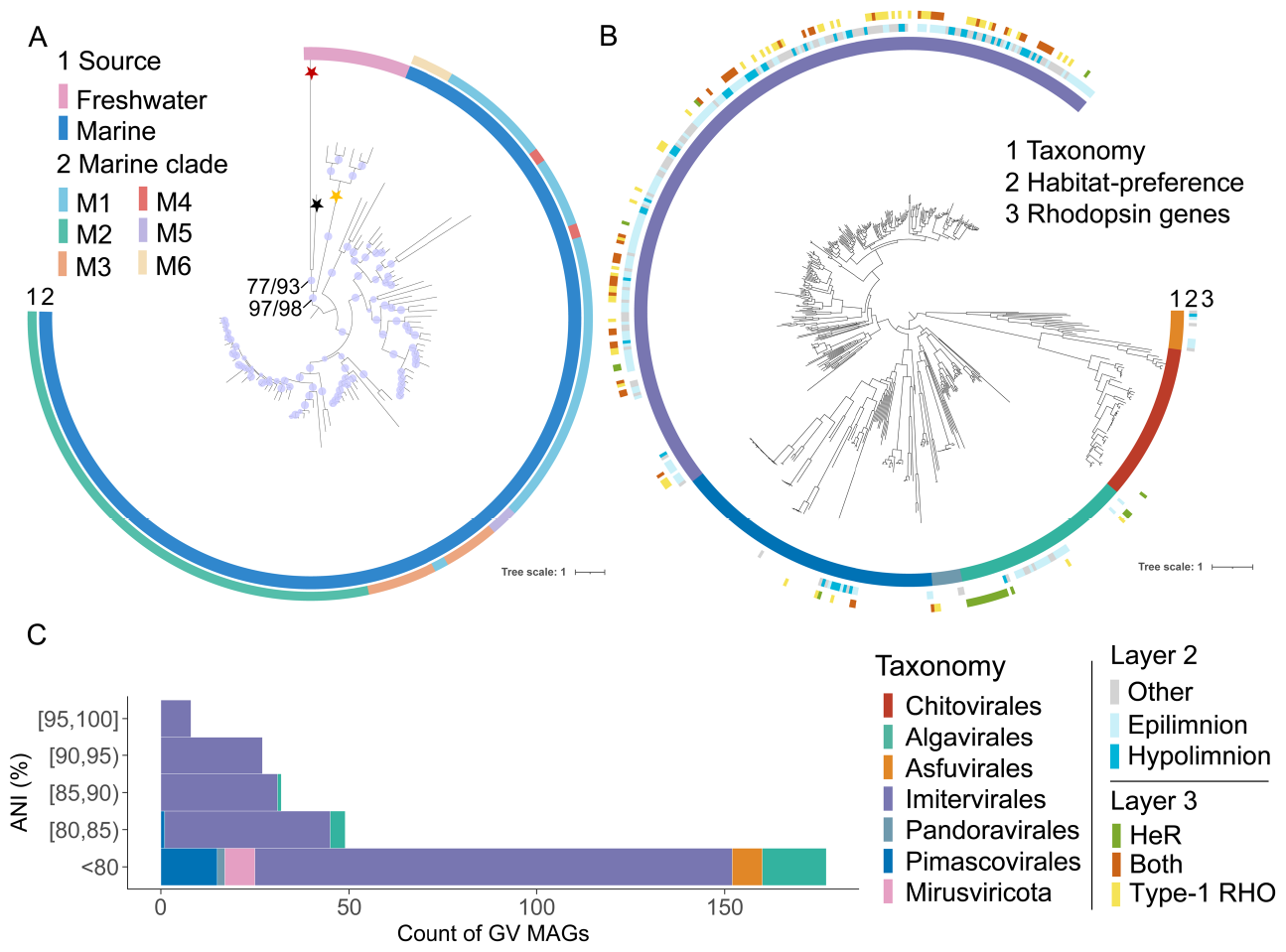
Phylogenetic diversity and novelty of GV MAGs. (A) Phylogenetic tree of the mirusvirus HK97 MCPs. Sequences from marine miruvirus MAGs [5] and freshwater miruvirus MAGs recovered in this study are included. The Ultrafast Bootstrap values are indicated by the sizes of the light purple circles on the branches. Certain key nodes indicating the divergences of different clades are indicated by values (aLRT/UFBoot) (see the Methods). Phylogenetic supports were considered high (aLRT ≥80 and UFBoot ≥95), medium (aLRT ≥80 or UFBoot ≥95), or low (aLRT <80 and UFBoot <95). The three proposed subclades of freshwater mirusviruses are marked with stars on the tree (subclade1: red; subclade 2: black; subclade 3: yellow). The tree is rooted between the freshwater and marine clades. (B) Phylogenetic tree of the recovered nucleocytoviruses. The tree was built using the concatenated protein sequences of seven genes (PolB, TFIIB, TopoII, A32, SFII, VLTF3, and RNAPL) and is rooted between the class *Pokkesviricetes* and *Megaviricetes*. The first outer layer of the tree (from the inside) indicates the taxonomic order of each GV genome, including our GV MAGs and the reference GV genomes. In the second layer, the uncolored branches of the tree are the reference genomes used to guide taxonomic assignment of our GV MAGs. “Other” means that the habitat preference was unclear. The third layer indicates the presence or absence of type-1 rhodopsin (Type-1 RHO)/heliorhodopsin (HeR)-encoding genes in our GV MAGs and the reference genomes. (C) The highest ANI value for each of our GV MAGs compared to the public GV genomes. Pair-wise ANIs were calculated between our MAGs and publicly available GV genomes and only the highest ANI for each MAG is plotted. ANIs <80% are not specified, being rather clustered into the “<80” category.

Following the screening approach guided by the mirusvirus HK97 MCP gene, we recovered eight mirusviruses from Lake Biwa, including the abovementioned circular genome (0074). HK97 MCP aside, other key components of the virion module were also shared with marine mirusviruses, including the genes encoding the portal protein, the capsid maturation protease, and the triplex capsid protein (Fig. S8A; Table S3). The phylogenetic trees inferred based on HK97 MCPs (Fig. 2B) and heliorhodopsins (Fig. S9) supported a distant evolutionary relationship between the newly identified freshwater clade and known marine mirusviruses. We further classified the freshwater clade into three subclades based on the topology of the HK97 MCP tree (subclade 1: 0074; subclade 2: 0010; subclade 3: the remaining six members). Consistent with the phylogenetic inferences suggested by the HK97 MCPs and heliorhodopsins, the GC content of freshwater mirusviruses was higher than that of marine mirusvirus clades (Fig. S8B). Further genomic analysis of this novel freshwater clade revealed that only 7.53– 22.87% of the genes per genome were homologous to those of the marine mirusvirus orthogroups (Table S3).

### Depth-specific distributions of GVs

The order-level community compositions and beta diversities among samples clearly revealed the succession of GV communities over the year (Fig. 3). After the water stratified (May–December), water circulation commenced and the GV communities of different water layers became well-mixed in February. A drastic shift in the hypolimnion community was apparent from January to February; however, the year-round cycle of the epilimnion community evidenced a more gradual change. This phenomenon is consistent with the mixing mechanism of lake water. As the temperature drops, the boundary of mixing water begins to gradually descend, and previously unmixed water is thus added to the epilimnion. In contrast, the hypolimnion is not affected until the mixing boundary attains the sampling depth (65 m). Throughout the stratification period, the compositional variation among communities in the hypolimnion was generally smaller than in the epilimnion. In line with NMDS analyses, most GV species (65.5%) were niche-specific. Specifically, 143 (48.8%) were epilimnion-specific and 49 (16.7%) were hypolimnion-specific.

**Figure 3.**
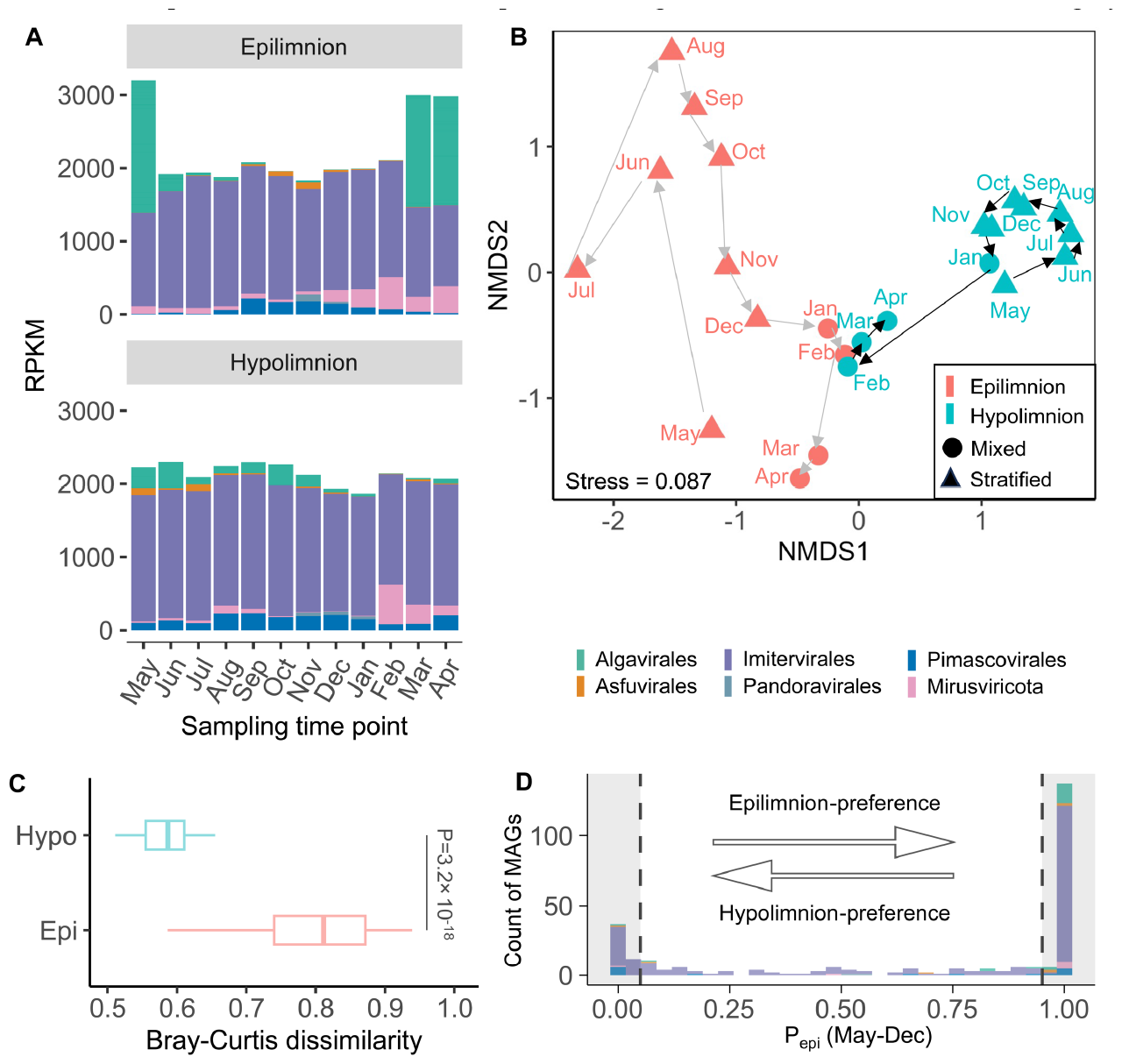
Spatiotemporal community shifts and vertical distributions. (A) Accumulative RPKM values across five orders of nucleocytoviruses and mirusviruses per sample. Within each bar, the abundance contributions made by each order or mirusvirus are indicated by different colors. Different blocks of the same color distinguish the MAGs. (B) NMDS plot of beta diversities across the samples. The pairwise beta diversities were derived by calculating Bray-Curtis dissimilarities based on the RPKMs from read mapping. Triangles refer to samples from the stratification period. Circles represent samples taken during water mixing. Hypolimnion samples are blue and epilimnion samples are red. (C) Comparison of the beta diversities of samples from the epilimnion and hypolimnion. The Bray-Curtis dissimilarities indicate the beta diversities for each pair of samples in the epilimnion or hypolimnion. (D) Habitat preference of each MAG as determined by the indicator P_epi_. As indicated in the gray background, MAGs with P_epi_ values >0.95 and <0.05 were defined as epilimnion- and hypolimnion-specific, respectively.

### Distinct dynamics of epilimnion and hypolimnion specialists

In general, GVs exhibiting different habitat preferences evidenced distinctive dynamic patterns. Epilimnion-specific GVs were typically opportunistic but hypolimnion specialists tended to be more persistent as indicated by their significantly higher persistence (*p*-value 1.6×10^-5^) during the water stratification period (Figs. 4, S10, and S11). Notably, nearly half (42.7%) of all epilimnion-specific GVs were present for only one month, whereas most (83.7%) hypolimnion-specific GVs were observed over at least two consecutive months.

**Figure 4.**
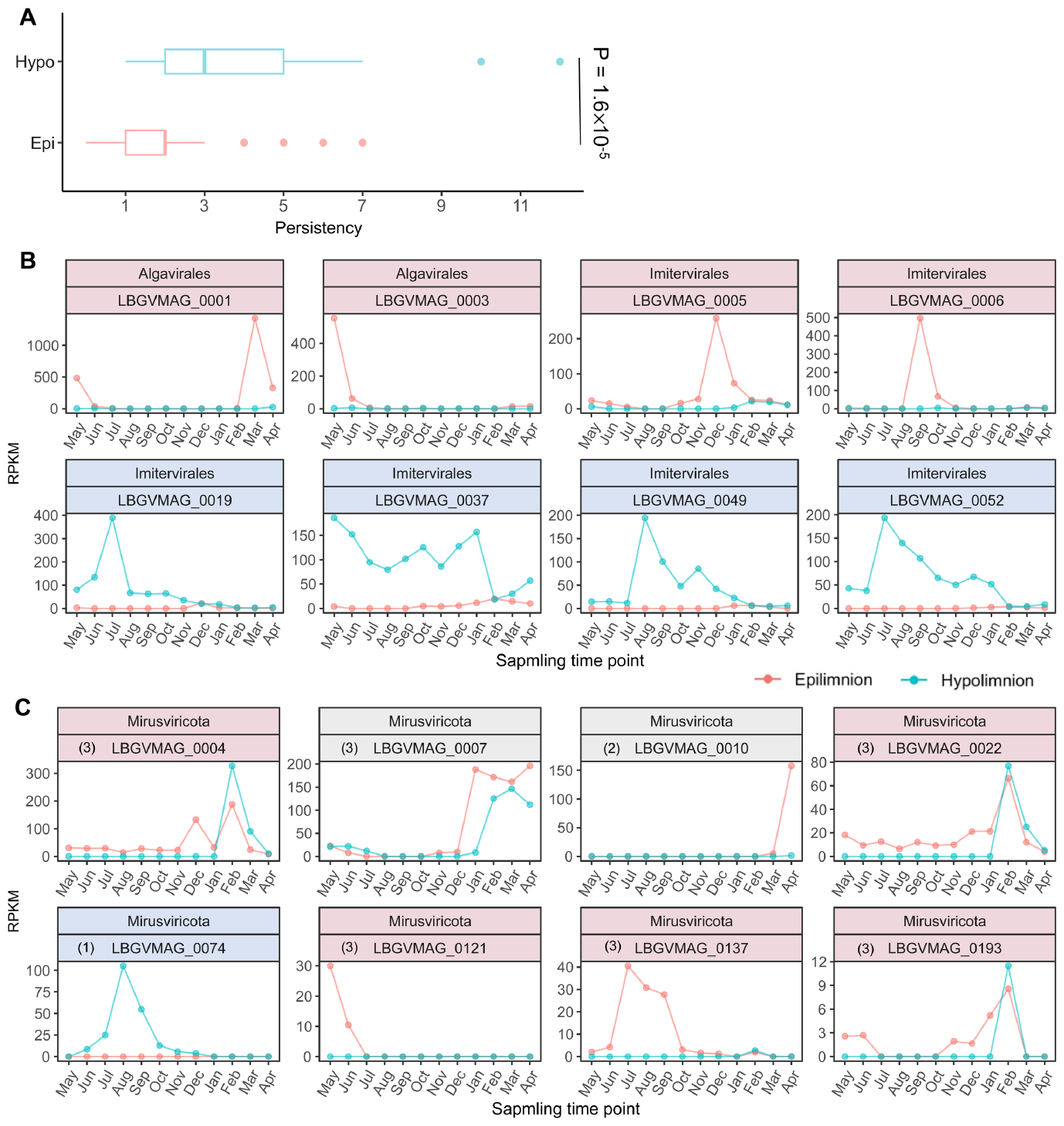
Community dynamics of GVs across water depths. (A) Persistence of GVs with different habitat preferences. Epi, epilimnion community; Hypo, hypolimnion community. (B) Community dynamics of the top four most-abundant epilimnion- and hypolimnion-specific nucleocytoviruses (determined by the maximum RPKMs across the samples). (C) Community dynamics of freshwater mirusviruses. Subclades of freshwater mirusviruses are indicated by the numbers in parentheses followed by the MAG IDs. In (B) and (C), the taxonomy and ID of each MAG are shown in the title of each box, and the color of the background indicates the habitat preference (pink, epilimnion-specific; blue, hypolimnion-specific).

Overall, imiterviruses dominated both water layers throughout the year, with the exception that four algaviruses collectively accounted for >50% of the relative epilimnion abundances in March, April, and May (Fig. 3A). Each of the four algaviruses (0001, 0002, 0003, and 0008) exhibited a relative abundance >5.3% for at least one month. Notably, 0001 accounted for 47.6% of the relative abundance in March. Among the four viruses, two (0001, 0003) were related to Yellowstone Lake phycodnaviruses (YSLPVs) of the AG_01 family, with ANIs of 87.5% and 84.1%, respectively [42]. The other two (0002, 0008) lacked any close relative in the database. GVs other than algaviruses predominated in the epilimnion in spring. The relative abundance levels of a mirusvirus (0007) ranked second and third among all GV MAGs in March (5.4%) and April (6.5%), respectively. Another mirusvirus (0010) ranked fourth in April, accounting for 5.3% of the relative abundance. Additionally, three viruses (0009, 0040, and 0018) with ANIs >87% to the reference genomes of the IM_01 family, and one virus (0014) with an ANI of 77.1% to a reference genome of the IM_12 family, were among the most abundant species identified. Each accounted for >2.1% of the relative abundance in the epilimnion during this period. Together, mirusviruses and nucleocytoviruses of the families AG_01, IM_01, and IM_12 were major players in the epilimnion of Lake Biwa in spring.

Furthermore, the dynamics of freshwater mirusviruses agreed with the differentiation of three subclades defined by the phylogeny of HK97 MCPs. The community dynamics of subclade 1 exhibited a typical hypolimnion-specific pattern (Fig. 4C), occurring only in the hypolimnion throughout the stratified period. In sharp contrast, subclade 2 was opportunistic, occurring exclusively in the epilimnion during April (at the end of water mixing) but absent from both water layers during the stratified period. The members of subclade 3 were typically epilimnion-specific and persistent.

## Discussion

### High-quality freshwater GV MAGs assembled from long reads

Although GVs exhibited high genetic diversity, they accounted for only ~2.0% of all metagenomic reads (Fig. S4), much lower than the value reported for prokaryotes in the same size fraction (0.2–5 μm) of the same samples (60.4%) [16]. This combination of high diversity and low read abundance posed a challenge for reconstructing GV MAGs. Nevertheless, our pipeline (Fig. S3) successfully captured such diversity in Lake Biwa as demonstrated by the inclusion of most viral *polB* genes in the MAGs (Fig. 1B). Although GV MAGs have been widely recovered using short-read sequencing [3–6], no long-read GV MAGs have previously been reported. Through comparisons with short-read MAGs, we validated the performance of long reads in terms of the recovery of more continuous and complete GV MAGs (Fig. S5).

Our spatiotemporal sample data focusing on an under-investigated freshwater system expand the known diversity of GVs, with many new species (97% of all species detected). A large proportion of MAGs had diverged from known reference species, primarily those with ocean-derived genomes [3–6]. Among those related to known GV references, most were closely related to freshwater-derived genomes, suggesting the existence of ecological barriers between the GVs of freshwater and marine environments. Furthermore, we discovered mirusviruses in the freshwater lake.

### A GV community specific to the dark hypolimnion

As observed for prokaryotes [16] and their viruses [43, 44], the GV community followed the physical structure of the seasonally stratified lake water column (Fig. 3B). Most species exhibited clear niche specificity for either the epilimnion or hypolimnion (Fig. 3D). The activities of hypolimnion-specific GVs were characterized by their persistent yet active turnover (Fig. 4A), associated with drastic increases and decreases in abundance over a short period of time. For example, the relative abundance of 0019 increased four-fold from June to July and then declined in August, indicating active virus replication and then decay in the hypolimnion (Figs. 4A and S11). Together with the observation that the viral host communities (microbial eukaryotes) are also vertically stratified [45], it is highly probable that hypolimnion-specific GVs infect hypolimnion-specific hosts. Unfortunately, our attempts to infer the hosts using gene phylogeny and genomic similarities to the isolated viruses were unsuccessful due to the lack of essential information in the database. Previous studies have revealed the existence of deep-water specific microbiomes including prokaryotes, their viruses, and eukaryotes [7– 10,45–50]. The persistence and exclusive presence of GV populations in the hypolimnion suggests that they are also part of a deep-water specific microbiome, likely playing important ecological roles in dark environments.

In contrast to hypolimnion-specific GVs, epilimnion-specific GVs were more diverse (Fig. 3D) and were typically opportunistic (Fig. 4A), in agreement with the observed associations between the GVs and opportunistic hosts including surface bloom-forming algae (Figs. 3A and S10) [51, 52]. GVs have been widely studied in the context of algal blooms, especially in the oceans, where they play critical roles in bloom termination [53–55]. However, the major GVs involved in freshwater spring blooms remain largely unknown. A recent study isolated a bloom-associated imitervirus of the IM_12 family from a freshwater lake [56], and another work characterized the GV community dynamics associated with algal blooms [11]. Our study revealed that mirusviruses and nucleocytoviruses of the families AG_01, IM_01, and IM_12 were among the most abundant GV lineages during the spring bloom of Lake Biwa.

A previous study reported high vertical connectivities among GV communities in marine environments that were suggested to be associated with sinking algae [57]. However, our data reveal a disconnect between the epilimnion and hypolimnion GV communities. Four algaviruses (0001, 0002, 0003, and 0008) that dominated in the epilimnion during the spring bloom (Fig. 3A) were rarely observed in the hypolimnion during the water stratification period (Fig. S10) despite the fact that significant sinking algal fluxes have been observed in Lake Biwa [58, 59]. The dominant algaviruses in the hypolimnion during lake stratification included one (0061) specific to the hypolimnion (Fig. S11) and another (0047) lacking a specific habitat preference (Fig. S12). These results suggest that the transportation of GVs from the epilimnion to hypolimnion associating with algal cells sinking is limited in Lake Biwa.

### Ubiquitous freshwater mirusviruses across water depths

GVs were previously known as large viruses of the phylum *Nucleocytoviricota*. Recently, an oceanic survey discovered a new GV phylum, *Mirusviricota* [5], which highlighted the importance of linking the evolutionary paths of the two viral realms *Varidnaviria* and *Duplodnaviria* [60]. The cited work reported that mirusviruses were abundant and widespread in sunlit oceanic areas. However, it remained unclear whether mirusviruses existed in freshwater ecosystems and aphotic regions. A recent work reported mirusvirus genomic fossils in the genomes of various eukaryotes including freshwater algae and cellular slime molds of soil, suggesting that their habitats are broad [61]. The recovery of mirusvirus genomes from Lake Biwa has firmly established their existence and activity in freshwater systems. The genomic features (Fig. S8) and phylogenetic inferences of the marker genes (Figs. 2A and S9) suggest that the freshwater mirusviruses form a distinct lineage that is placed outside the marine clade, and that the unique gene repertoires indicat an untapped source of genetic diversity.

We found one circular mirusvirus genome (0074) confined to the dark hypolimnion (Fig. 4C), possibly infecting hypolimnion-specific hosts. A previous oceanic survey of GVs across water columns from 2 to 5,500 m did not detect any mirusvirus genome below the photic layer [6]. Recent studies have reported that the algae *Aurantiochytrium limacinum* (Labyrinthulea) [62] and *Cymbomonas tetramitiformis* (green algae) [63] are highly probable hosts of mirusviruses. However, these organisms typically inhabit sunlit regions. Our observation of a hypolimnion-specific mirusvirus highlights the potential roles of mirusviruses as components of the deep-water specific microbiomes of freshwater ecosystems.

### GV rhodopsins in a dark environment

Our results also suggest that GV rhodopsins (heliorhodopsins and type-1 rhodopsins) might have light-independent functions. The viral type-1 rhodopsins of nucleocytoviruses [64– 69] were previously thought to be involved in light absorption and sensing, in turn influencing the behaviors of photosynthetic hosts during infection. Heliorhodopsins are actively expressed in marine mirusviruses, especially those of sunlit oceans [5]. In the present study, 15 (30.6%) hypolimnion-specific and 57 (38.5%) epilimnion-specific GV species harbored rhodopsin genes (Fig. S13). Notably, heliorhodopsin genes were present in most freshwater mirusvirus MAGs (7/8) recovered in this study. A recent work found that all heliorhodopsins in theionarchaeal (archaeal) genomes were identified in light-insufficient environments [70]. Our report of rhodopsin genes among hypolimnion-specific GVs point to previously under-investigated functions in dark habitats.

### High dispersal rates of GVs

We recovered three imiterviruses (Fig. S7) from Lake Biwa that were nearly identical (ANIs of 99.2%, 98.2%, and 98.2%, respectively) to those of Lake Lanier in North America [4], suggesting a recent dispersal event between two lakes ~11,220 km apart. In terms of microbial dispersal, prokaryotes have been the foci of previous studies. However, a vigorous debate continues regarding whether these microbes are globally distributed at the species level [71–74]. Compared to prokaryotes, viruses generally exhibit higher mutation rates [75], suggesting that global distribution of the same GV species is attributable to a high dispersal rate rather than slow local diversification. Although GV dispersal has not been well-studied, a few long-distance dispersal events across continents have been suggested [76]. Two mimiviruses with nearly identical genomes (ANI of ~99.9%) have been isolated in Japan [77] and the UK [78]. Two marseilleviruses sharing ~98.5% of ANIs have been isolated in China (GenBank MG827395) and France [79]. Although these observations imply long-distance GV dispersal, a recent study revealed a high degree of endemism among lakes of the two poles and within each polar region [80]. Quantitative studies on the dispersal vs. local diversification rates of GVs will aid further understanding of the processes shaping the global biogeography of GVs. Deep lakes, given their high levels of physical isolation, would serve as useful models for such quantitative studies.

## Conclusion

Using a combination of spatiotemporal sampling and long-read metagenomics, we reveal the previously overlooked diversity of freshwater GVs, as evidenced by 285 new species and the discovery of viruses of the phylum *Mirusviricota* in a freshwater lake. We demonstrate the habitat preferences and community dynamics of GVs. Most nucleocytoviruses and mirusviruses could be clearly classified as being specific to either the epilimnion or hypolimnion. Epilimnion specialists tended to be opportunistic, while hypolimnion specialists were typically more persistent. These distinctive dynamic patterns suggest that GVs employ diverse ecological strategies, and our work paves the way toward a better understanding of the roles played by GVs in microbial ecosystems. Specifically, the strong seasonality in the epilimnion suggests that GVs make significant contributions to the plankton community shift in the lake. Not only nucleocytoviruses but also mirusviruses are major players during the spring bloom. Conversely, persistent hypolimnion specialists that nonetheless exhibit active turnover suggest that GVs also play unique and important roles in the hypolimnion-specific microbial ecosystem. Furthermore, our observation of nearly identical GVs in lakes of different continents further suggests a ubiquitous distribution of GVs at the species level, highlighting the role of dispersal in shaping the global distributions of GV communities. In light of this, we call for research on GV host interactions, diversity, and biogeography in freshwater lakes worldwide, providing key insights into the eco-evolution of GVs in such unique ecosystems.

## Supporting information

Legends and titles of supplementary figures 1-13 and tables 1-3; Supplementary Methods

## Acknowledgments

This study was supported by the Center for Ecological Research, Kyoto University; the Joint Usage/Research Center, the Kyoto University Foundation; a JST FOREST Program grant JPMJFR2273; and JSPS KAKENHI grants 22H00384, 16H06279 (PAGS), 18J00300, 22H00382, and 22K15182. All computations were done on the Supercomputer of the Institute for Chemical Research, Kyoto University.

## Data availability

The nucleotide and protein sequences of the LBGVMAGs generated in the present study are available from https://www.genome.jp/ftp/db/community/LBGVMAGs/ at GenomeNet.

