## Supplementary material for "Spatiotemporal dynamics revealed the dark water community of giant virus from a deep freshwater lake": Legends and titles of supplementary figures 1-13 and tables 1-3; Supplementary Methods

### Supplementary Information

#### Supplementary Figures

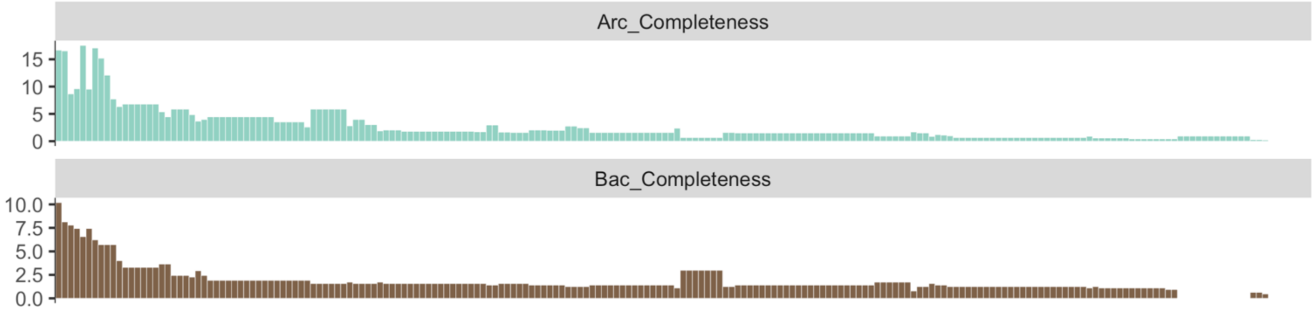


**Figure S1. Completeness value of 207 nucleocytovirus reference genomes estimated by CheckM.** Each bar represents one genome. The upper panel was estimated using a set of bacterial marker genes, and the lower panel used a set of archaea marker genes.

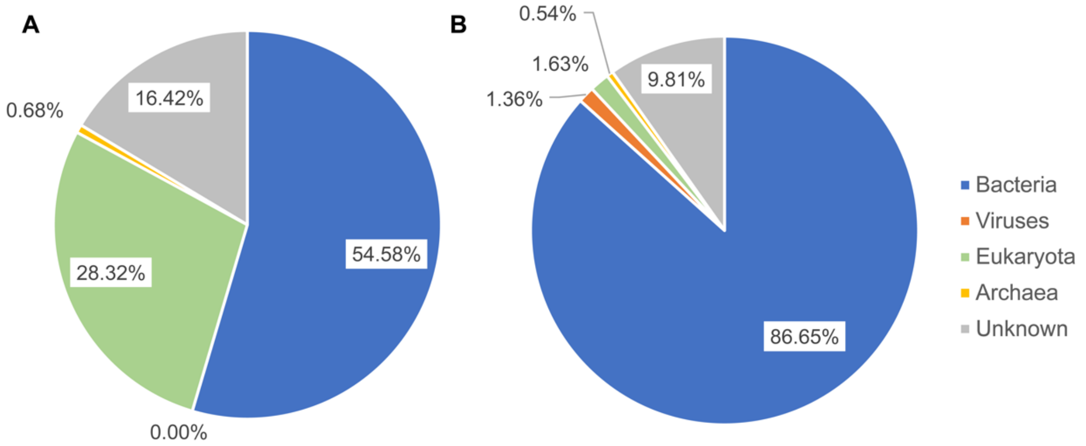


**Figure S2. Taxonomic annotation of nonviral contigs and partial regions within the chimeric contigs identified.** We assigned taxonomy to contaminated (A) contigs and (B) regions removed.

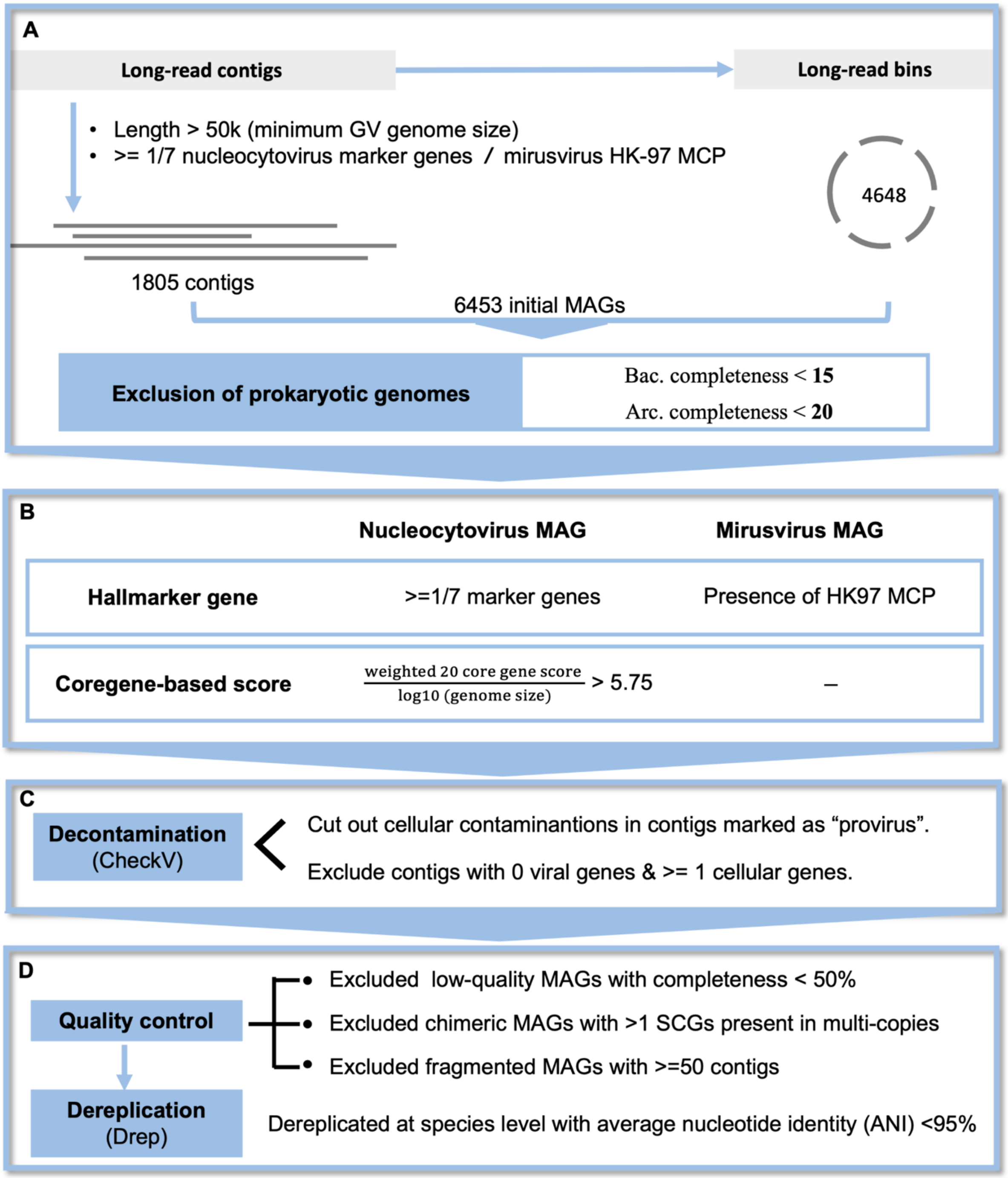


**Figure S3. Pipeline for recovering long-read GV MAGs**. In this pipeline, we not only recovered bins but also recruited nearly complete GV contigs by taking advantage of long-read sequencing. A customized pipeline guided by MCP gene was used to recover miruvirus MAGs.

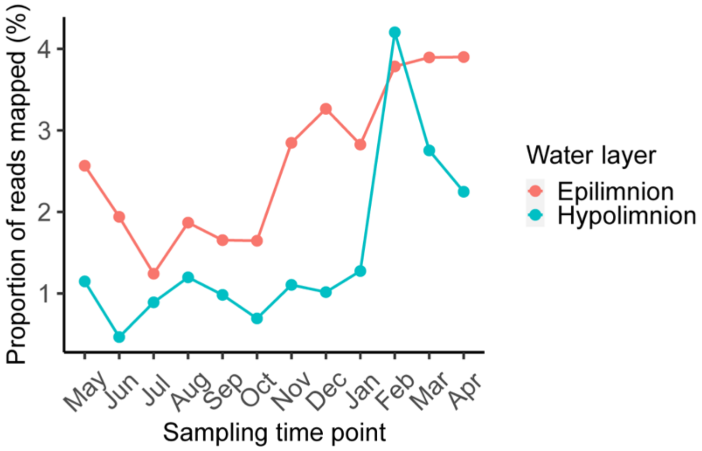


**Figure S4. Proportion of short reads mapped onto GV MAGs across 24 samples.** The short reads were mapped to the GV MAGs, and the proportion of mapped reads in each sample was plotted.

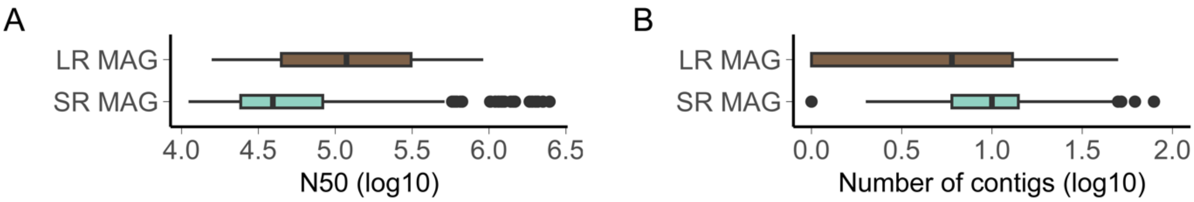


**Figure S5. Comparison between long-read and short-read MAGs.** We compared (A) the N50 and (B) number of contigs per MAG between our long-read GV MAGs (LR MAG) and high/medium-quality short-read MAGs from GVDB (SR MAG).

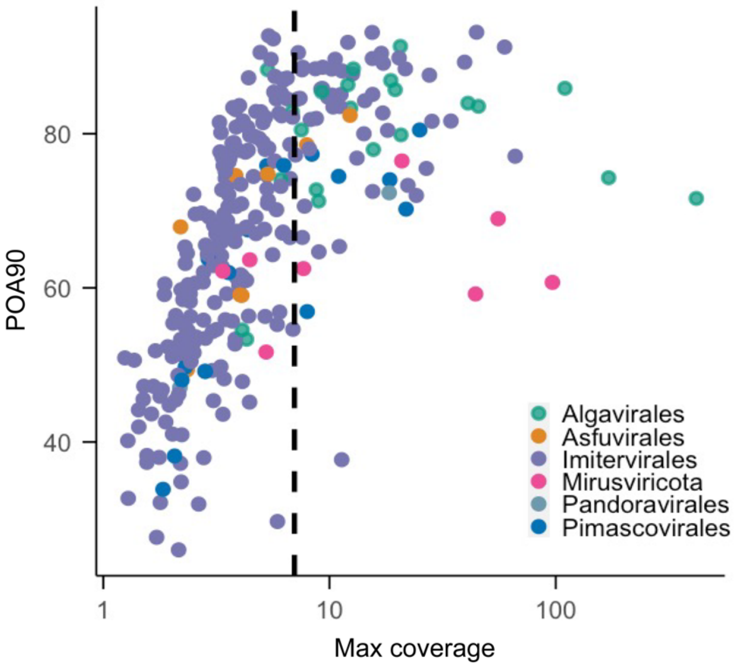


**Figure S6. POA90 score of GV MAGs.** The dashed line indicated the threshold of coverage (7x), suggesting the indel errors were effectively corrected in the GV MAGs.

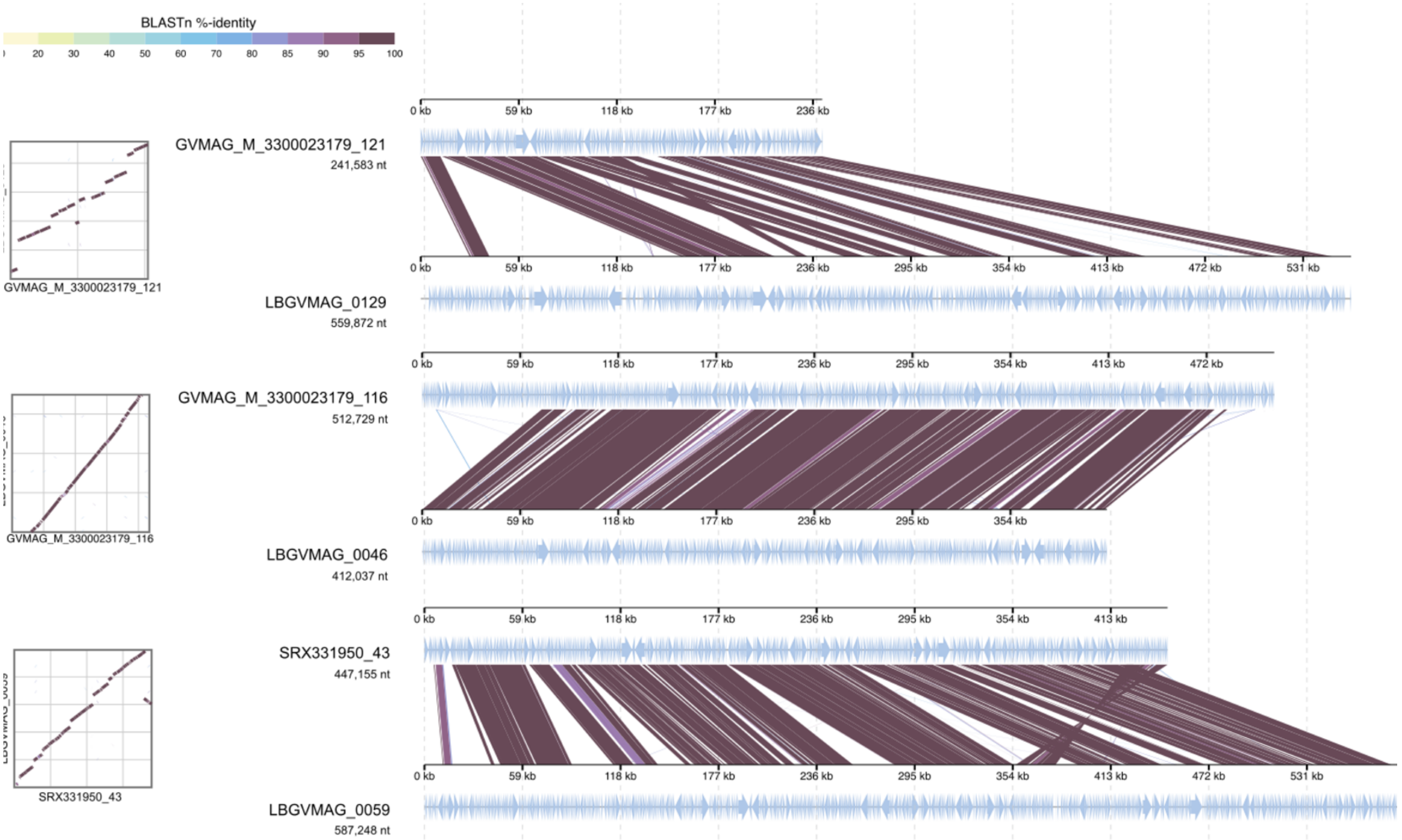


**Figure S7. Blastn alignment of almost identical GV genome pairs.** From the top to bottom, the average nucleotide identities (ANIs) of the pairs were 99.18%, 98.21%, and 98.19%, respectively. Within each pair, the MAG on top was from Lake Lanier, and the bottom was GV MAG recovered in this study from Lake Biwa. Contigs were reordered and concatenated within each genome for better visualization.

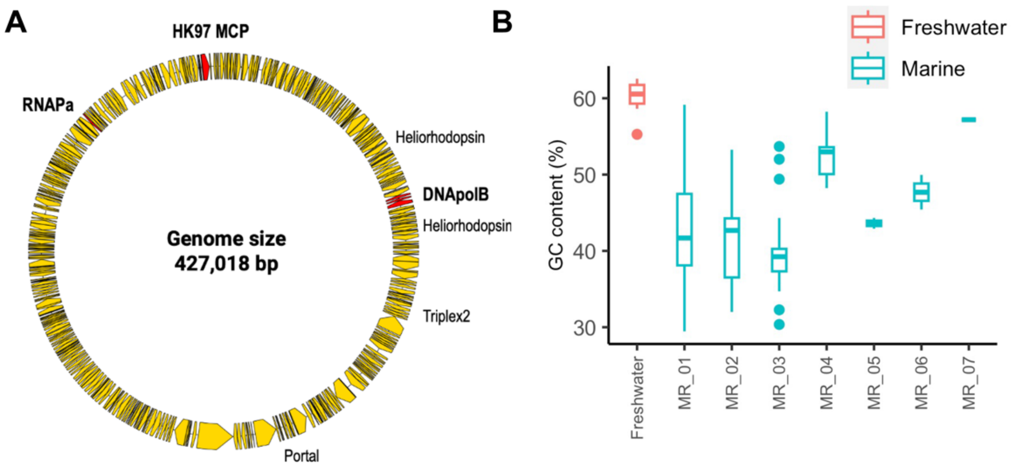


**Figure S8. Comparison of genomic features between freshwater and marine mirusviruses.** (A) We visualized key homologies of marine mirusviruses found in a freshwater mirusvirus genome (0010) as an example. Note that the genome was shown in circular only for the convenience of visualization. (B) Comparison of GC content between freshwater and marine mirusvirus clades (MR_01-07).

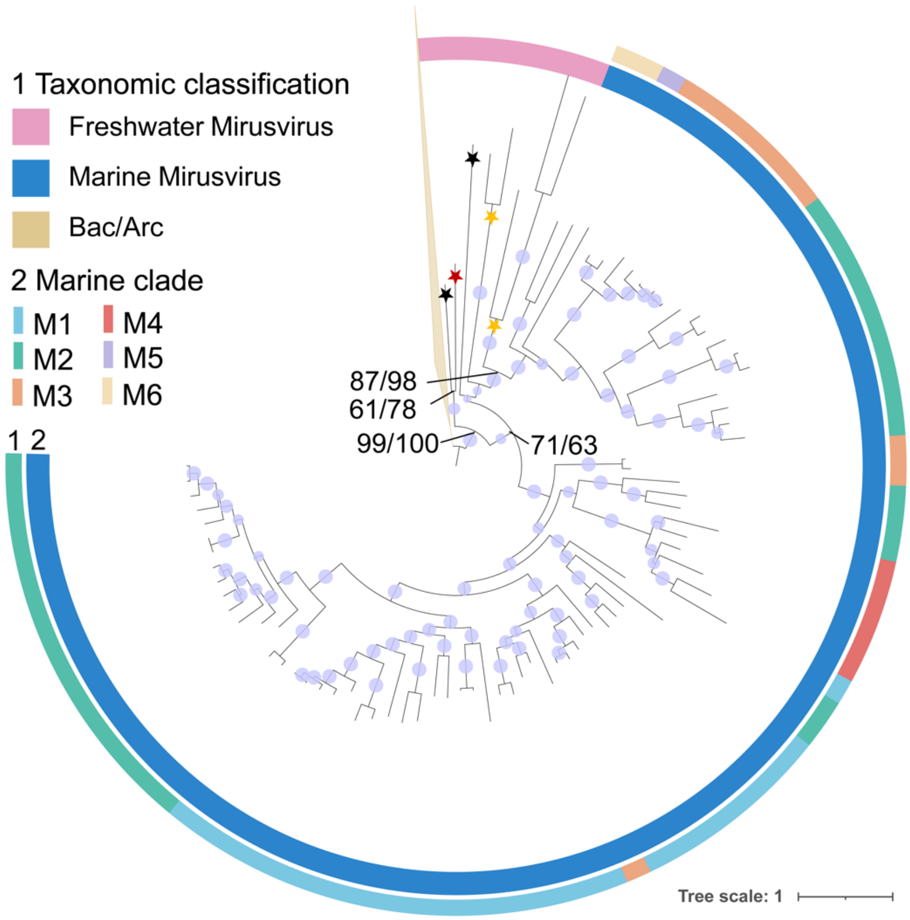


**Figure S9. Phylogenetic tree of mirusvirus heliorhodopsins**.

The tree included sequences of freshwater mirusviruses, marine mirusviruses and reference sequences of bacteria and archaea downloaded from RefSeq database. The tree was rooted between the clasped branch of bacterial and archaea heliorhodopsin sequences and the rest mirusvirus sequences. Three proposed subclades of freshwater mirusviruses were marked with stars on the tree (sub-clade 1: red; sub-clade 2: black; sub-clade 3: yellow). The Ultrafast Bootstrap value of each node was shown by the size of circles on the nodes. Certain key nodes indicating the divergence of different clades were specified by Ultrafast Bootstrap values (aLRT/UFBoot) (see the Methods). Phylogenetic supports were considered high (aLRT>=80 and UFBoot>=95), medium (aLRT>=80 or UFBoot>=95) or low (aLRT<80 and UFBoot<95).

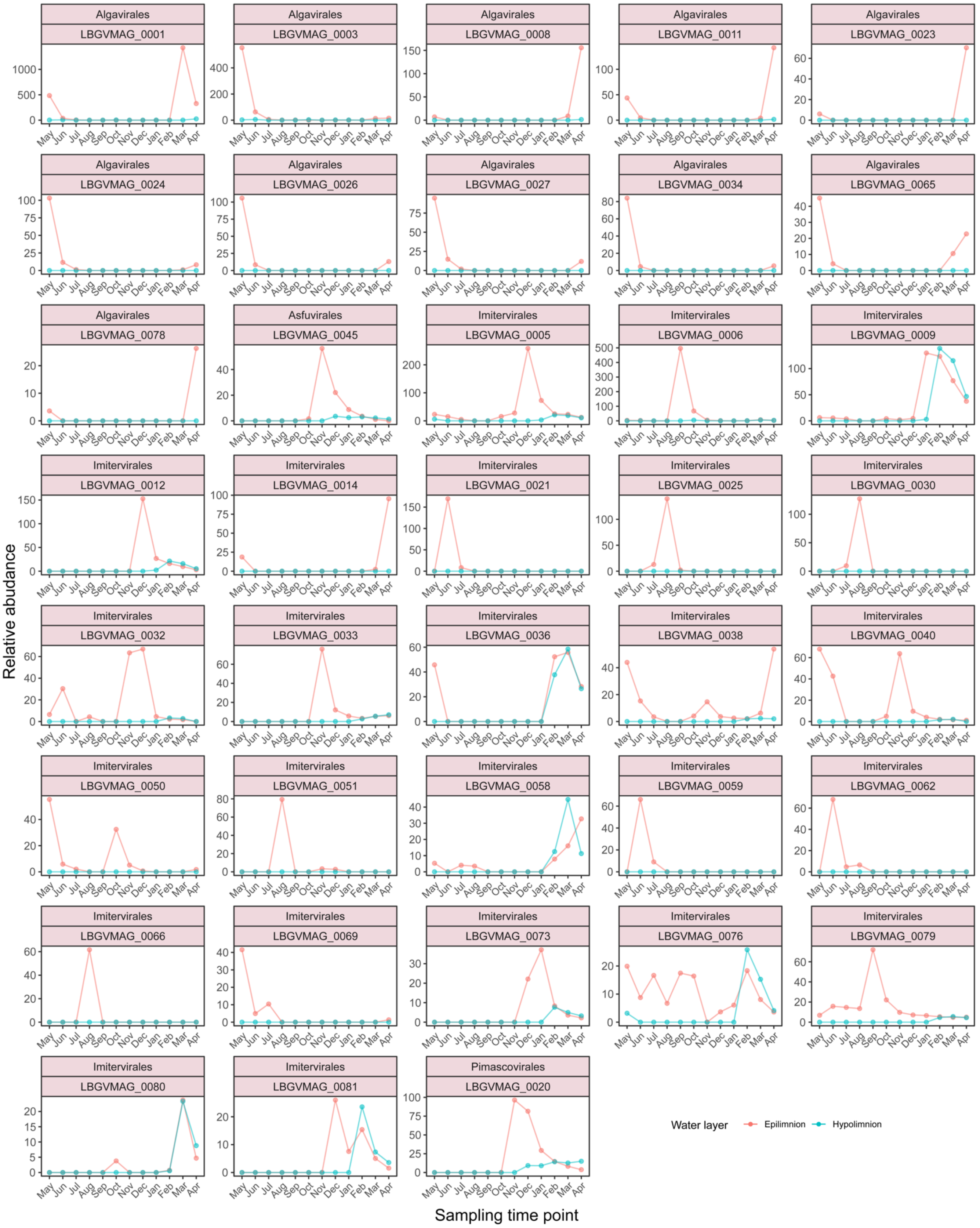


**Figure S10. Community dynamics of epilimnion-specific nucleocytoviruses.** Epilimnion-specific MAGs with coverages >7× were selected for this analysis based on relative abundance (RPKM). The title of each box included taxonomy (the first line) and MAG ID (the second line).

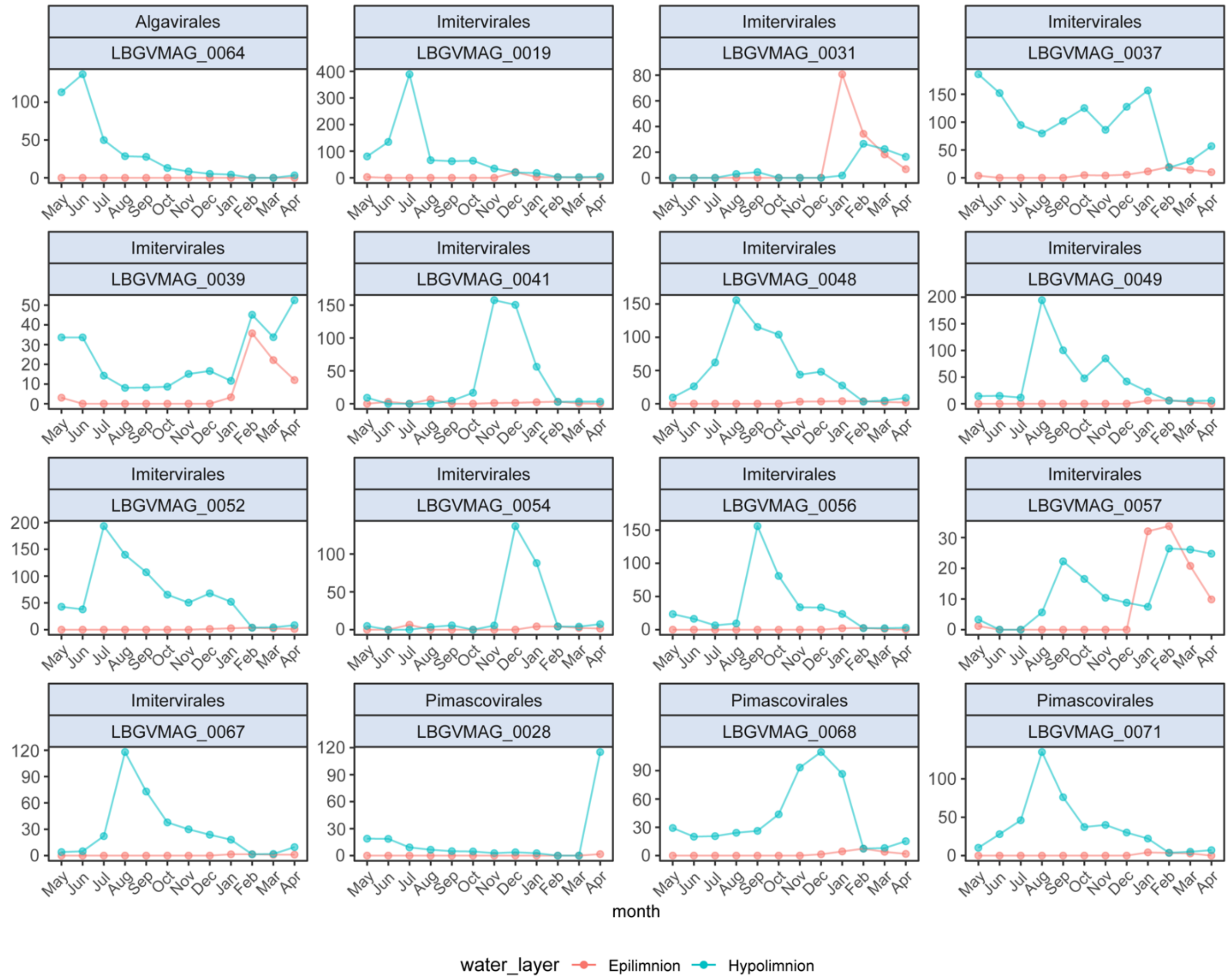


**Figure S11. Community dynamics of hypolimnion-specific nucleocytoviruses.** Those hypolimnion-specific MAGs with coverages > 7× were selected for this analysis based on relative abundance (RPKM). The title of each box included taxonomy (the first line) and MAG ID (the second line).

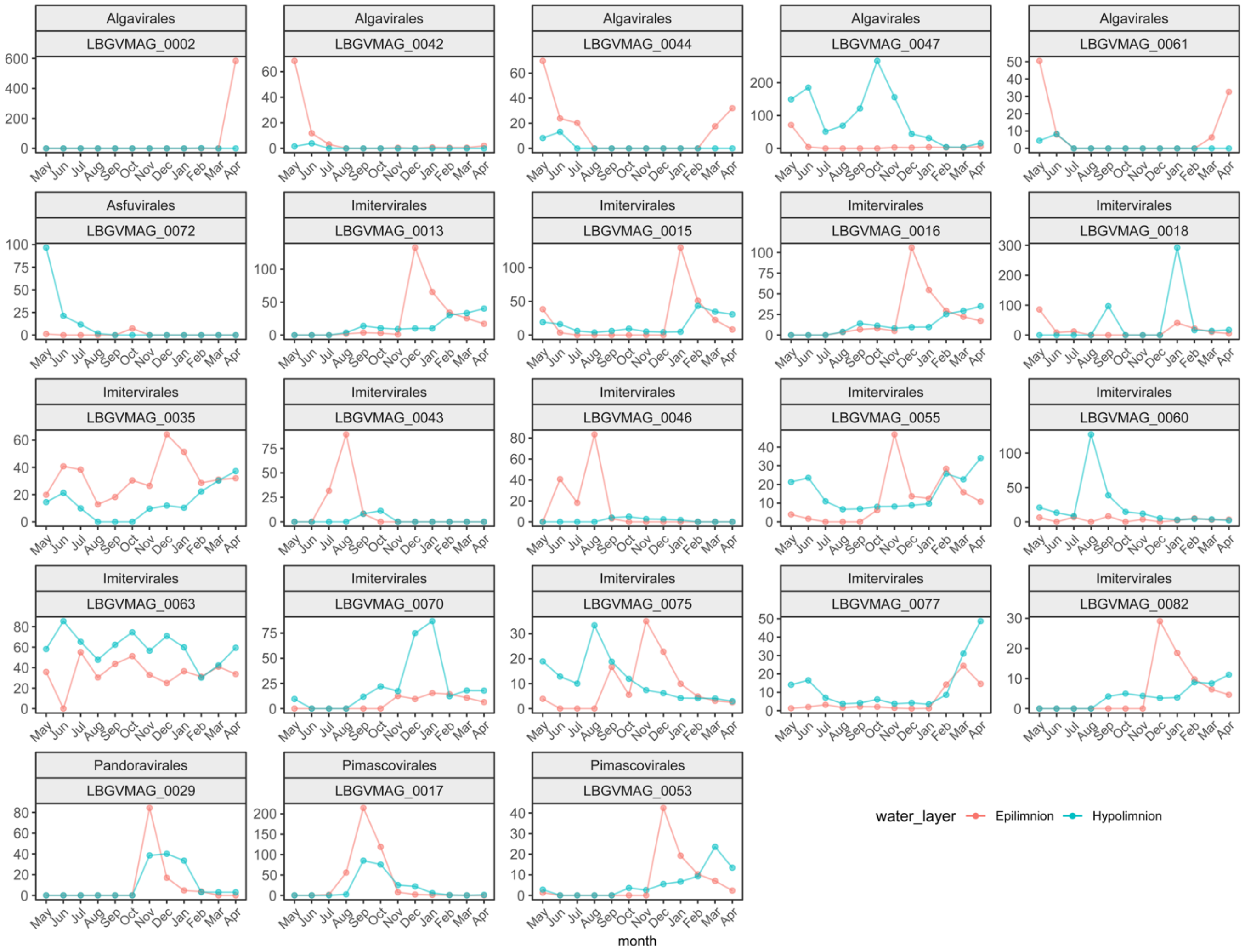


**Figure S12. Community dynamics of nucleocytoviruses with no identified habitat preference.** Those vertically ubiquitous MAGs with coverages > 7× were selected for this analysis based on relative abundance (RPKM). The title of each box included taxonomy (the first line) and MAG ID (the second line).

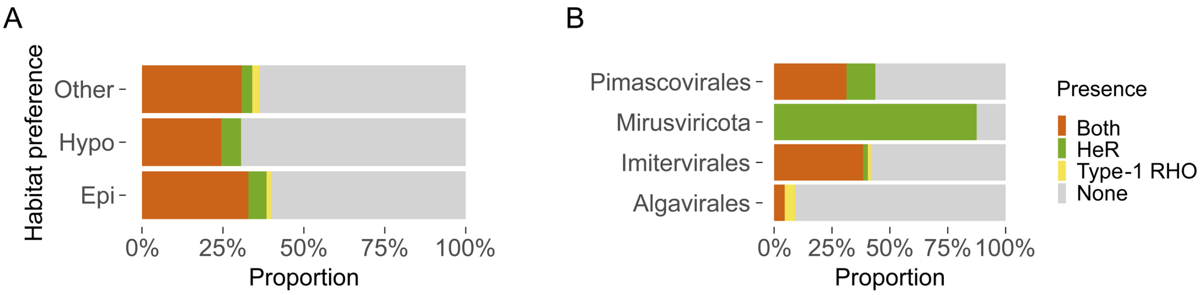


**Figure S13. Microbial rhodopsin-encoding gene detected from GV MAGs.** In both figures, we focused on the presence or absence of rhodopsin genes in the MAGs. Multiple copies of a type-1 rhodopsin (Type-1 RHO) or heliorhodopsin gene (HeR) within a MAG were treated equivalently to single copy, and thus marked as 'present'. The proportion of MAGs with rhodopsin genes present was calculated as the percentage of all GV MAGs in which rhodopsins were identified.

#### Supplementary Tables and Legends

**Supplementary Table 1. Description of 293 LBGVMAGs.** This table includes basic descriptions of the 293 GV MAGs recovered in this study, including the genome size, number of contigs, N50, habitat preference, number of each marker gene detected, abundance profiles, etc..

**Supplementary Table 2. ANI values calculated for GV MAGs.** Pair-wise ANI comparisons were done for each GV MAGs against the custom database in this study (See the Methods). only the pair with the highest ANI value for each GV MAG was selected and plotted, thereby highlighting the closest genomic relationships with GV genomes from other environments. The sources of the GV genomes from other enviroments are also specified in this table. GV MAGs without specific ANI values, due to large divergence from GV genomes in the custom database, are not included in this table.

**Supplementary Table 3. Shared orthogroups of marine and freshwater mirusviruses.** This table lists all orthogroups from marine mirusviruses that are present in at least one freshwater mirusvirus MAG recovered in this study. Gene annotations by three public databases (See the Supplementary Methods) are included.

#### Supplementary Methods

##### Exclusion of prokaryote-like MAGs

As GVs accounted for a minor proportion of the assembled contigs and bins compared to prokaryotes, we excluded prokaryotic sequences before downstream screening. We employed CheckM v1.2.2 [1] to estimate the completeness of each GV MAG candidate as a bacterial or archaeal genome. To minimize false positives, we tested the threshold for the completeness score to exclude a GV MAG using 207 reference GV genomes [2] manually collected from RefSeq database and related publications (Fig. S2). A completeness score of 15 for bacteria and 20 for archaea were determined as balanced thresholds to exclude prokaryote-like genomes. GV MAG candidates with completeness scores higher than either of these two thresholds were excluded.

##### Screening for GV MAGs

For nucleocytoviruses, the weighted scoring method was determined from the presence of 20 core genes [2]. Gene predictions of the GV MAG candidates were done using Prodigal v2.6.3 with "-meta" mode [3]. Sequences of these 20 core genes, universal or nearly universal in all *Nucleocytoviricota* lineages, were downloaded from the Nucleo-Cytoplasmic Virus Orthologous Groups (NCVOGs) database [4]. Multiple alignments for each NCVOG were performed using MAFFT v7.505, then the corresponding hidden Markov model (HMM) was built with the function ‘hmmbuild’ of HMMER3 v3.4 [5] and used to determine the presence or absence of 20 nucleocytivirus core genes in each GV MAG candidate. Then the weighted score of each MAG was calculated as below:

$$Core\ gene\ index = \frac{\sum_{k=1}^{20} Weight_k * X_k}{\log_{10}(genome\_size) - 4}$$

where $Weight_k$ was given based on prevalence level of core gene K among the 11 families of given reference genomes (e.g., if gene K is present in all 11 out of 11 families, $Weight_k = 1.1$) and $X_k$ is a binary variable representing whether the core gene K is absent (0) or present (1) in the MAG. Putative nucleocytovirus MAGs were selected with a cut-off of 5.75 [2]. Finally, we again curated the resulting MAGs by selecting those with at least one of the seven marker genes described above [6]. Occurrences of the seven marker genes were determined using the tool “ncldv_markersearch” [7].

For the detection of mirusviruses, we screened for the mirusvirus marker gene, HK-97 MCP gene, in the MAGs. Protein sequences of the HK-97 MCP gene from marine mirusvirus MAGs published recently [8] were collected for generating the HMM model as described above, which was used for screening for mirusvirus MAGs.

**Removal of cellular contamination from GV MAGs**

Since we recovered GV MAGs from the same size fraction (0.2–5 μm) as prokaryotes, mis-binning of a prokaryotic contig was the major source of contamination. Therefore, we further removed prokaryotic sequences from the remaining GV MAGs by screening each contig with “end_to_end” pipeline of CheckV v1.0.1 (database v1.4) [9]. When CheckV detected at least one microbial gene but no viral-specific gene from a contig, the contig was regarded as prokaryotic and consequently removed from the MAG. Through this process, we further found a few contigs annotated as “provirus” by CheckV. Manual inspections indicated that these contigs were not *bona fide* proviruses (GV genomes endogenized in the host genome) but likely resulted from mis-assembly of GV and prokaryotic genomes. For such contigs, we cut out the segments assigned as “host region”, which were defined as non-viral due to the dominance of cellular signals by CheckV. For all regions and contigs removed through this process, we additionally performed taxonomic annotations using CAT [10] to ensure that the removed regions and contigs were indeed contamination (Fig. S3). In this way, we thoroughly removed cellular contamination from the GV MAGs.

**Exclusion of chimeric, low-quality, and fragmented GV MAGs**

We removed chimeric MAGs in which more than one single-copy marker gene was found in multiple copies [7]. We also excluded fragmented MAGs with more than 50 contigs from further analysis [11]. Additionally, we removed low-quality MAGs that showed the completeness score lower than 50 [12] using CheckV v1.0.1 [9]. In the case of multi-contig MAGs, we concatenated all the contigs into a single sequence and subsequently performed CheckV on the concatenated contig as CheckV only accepts single-contig input.

**Dereplication of GV MAGs**

The redundancy of GV MAGs generated from the above processes has been removed based on their average nucleotide identity (ANI) using dRep v.3.2.2 with "--S_algorithm ANImf --clusterAlg single -sa 0.95" parameters. Also, the completeness and contamination scores generated by CheckV v1.0.1 was given by "–genomeInfo" parameter [13]. ANImf with a minimal ANI threshold of 95% was chosen for the secondary clustering because 95% ANI is a commonly used species boundary for GVs [6].

**Calling polBs from contigs**

We called *polBs* from all the assembled contigs following a pipeline proposed in a previous study [14]. Only *polB* sequences with >500 codons and >7× contig read coverage were used for the analysis. We clustered *polB* nucleotide sequences with cd-hit v4.8.1 [15] at 98% identity and performed the alignment against contigs in GV MAGs.

**Detection of type-1 rhodopsin gene**

We identified rhodopsins by hmmsearch program of HMMER v3.4 against the Pfam model for type1 rhodopsins (PF01036) with a threshold E-value of $1\times10^{-3}$.

**Indel error assessment of long-read GV MAGs**

The indel errors are common in long-read assembly [16] due to the higher nucleotide error rate than the short-read platforms. Even though the contigs have already been error-corrected (polished) using the long and short read mapping, some errors remained inevitably [17]. Following the previous study [17], POA90 score was defined as the proportion of amino acid sequences in which >90% of the length was aligned to a UniRef90 sequence.

**Comparisons of genome contents between freshwater and marine mirusviruses**

Orthogroups of marine-originated mirusviruses were generated by OrthoFinder v2.5.5 with default parameters [18]. Orthogroups involving more than 10 mirusvirus genomes have been retained. Annotations of orthogroups were done based on Pfam [19], PDB [20], and SCOP70 database [21] by hhsuite v3.3.0 [22] and hmm files were generated for each orthogroup with hmmbuild program of HMMER v3.4. Then, genes of all freshwater mirusvirus genomes were predicted using prodigal v2.6.3 with "-meta" mode [3] and searched against the marine mirusvirus orthogroups by hmmsearch program of HMMER v3.4 with a threshold E-value of $1\times10^{-3}$.
